# Population modelling insights of extinct environments: the case of the Kem Kem palaeocommunity

**DOI:** 10.1101/2021.09.07.459352

**Authors:** Lucas dos Anjos

## Abstract

The Kem Kem beds are well-known palaeontological deposits. Among the species that lived there, there are some large theropods, such as *Deltadromeus agilis, Carcharodontosaurus saharicus*, and *Spinosaurus aegyptiacus*. It is possible that these large predators were facultative scavengers, and they could compete for carrion. In the present paper, I simulate a small community module of this environment, consisting of Carrion, Fishes, *Spinosaurus*, and a functional group composed of large terrestrial Theropods. I assume that these top predators feed on carrion, but they also have exclusive food sources. I show that these exclusive food sources could have assured the possibility of coexistence, and in their absence, one top predator could be locally extinct.

## 1 Introduction

The Kem Kem deposits in Morocco are a rich palaeocommunity composed of diverse groups [1]. One peculiar feature of this palaeocommunity is that palaeontologists find more fossils of large-predators dinosaurs than the corresponding herbivores, which led some authors to name this patter as “Stomer’s riddle” [1, 2]. Some authors have suggested that the reasoning behind this pattern is a collecting bias [2, 3]. However, other authors suggest that the pattern can be indeed a natural phenomenon in opposition to collecting bias [1]. Assuming that the pattern is caused by actual biological processes (i.e., not a collecting artefact), one potential explanatory mechanism is resource partitioning among large predators [4, 5], which is evidenced by calcium isotopes [6]. Among the coexisting large-bodied theropod predators of this environment, there are, for instance, *Deltadromeus agilis* Sereno et al., 1996 [7], *Carcharodontosaurus saharicus* Stromer, 1931, and *Spinosaurus aegyptiacus* Stromer, 1915 sensu Smyth et al., 2020 [8].

In extant communities, as the African savannah, large predators interact with each other in several ways. These interactions can be fulfilled by direct contact or, likely more commonly, indirect dispute through scents and display [9]. The most evident direct interaction is the interspecific competition, which in fact occurs for some pairs of species [10]. Another important interaction is the intraguild predation, which involves both interspecific competition and predation among the involved species, and this type of interaction also potentially occurs in African savannah [9]. Regarding indirect disputes, a top predator can inhibit the foraging behaviour of mesopredators or shift their foraging range [11, 12, 13]. This latter mechanism of avoidance is of particular interest for the Kem Kem palaeocommunity, since it does not necessarily involve actual clash, something that to my knowledge was not yet found in the fossil record among the large predators.

In this paper, I study a resource competition system between top predators of the Kem Kem Group from the Cretaceous period. I assume that much like modern African predators, the top predators of that time also were opportunistic scavengers, and also displayed resource competition for carrion. I also assume these top predators had indirect contact displaying density-mediated interactions [14]. Because scientists can not directly observe the dynamics of extinct communities, they have an extra complication compared with extant communities. In this circumstance, mathematical models of population dynamics may be useful tools for palaeoecology due to the scarcity of data these systems possess [15, 16]. Within this framework, the objective of this work is to answer the following questions: (i) how does the density-mediated interaction influence the coexistence of two top predators living in the same environment?; (ii) how does an increase in food input of one top predator affect its competitor?; and (iii) how does a variation in the shared resource can affect the top predators’ densities?

## 2 Methods

### 2.1 System description

The interactions studied in this paper are presented in Figure 1. The analysed system is a fragment of the potential palaeocommunity modules of that ecosystem, consisting of two top predators, *Spinosaurus* (*S*) and other Theropods (*T*); “Fishes” (*F*) that represent the community of fish species that are consumed by *Spinosaurus*; “Sauropods and others” that are items consumed by other Theropods. Carrion dynamics have an important role in extant ecosystems [17, 18, 19], and could also be important in the Kem Kem palaeoenvironment. Given that, I assume that the two top predators compete by resource competition for the Carrion (*C*), and may influence each other through a density-mediated interaction (DMI, hereinafter) regarding this consumption.

**Figure 1:**
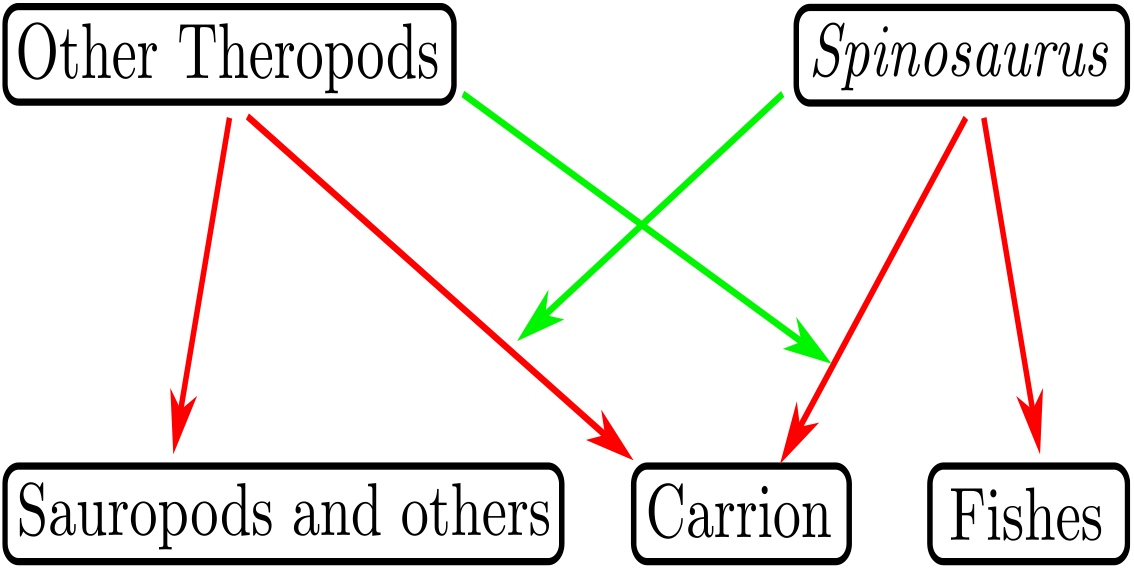
Hypothetical interactions of the Kem Kem palaeoenvironment studied in the present work. Red arrows indicate consumption, while green arrows are the top predators’ density-mediated interaction (DMI).

#### 2.1.1 Environment

The location of the study system is named the Kem Kem Group in Morocco [1]. This palaeoenvironment is characterized by having several microhabitats, which could provide opportunities for niche diversification [20, 21]. The vertebrate fauna is represented by elasmobranchs, osteichthyes, sarcopterygians, amphibians, lepidosauromorphs, turtles, crocodyli-forms, pterosaurs, non-avian dinosaurs, and possible others [1] (and references therein).

#### 2.1.2 Spinosaurus

It has been showed that *Spinosaurus* had anatomical adaptations to pursue [22] and catch fish [23, 24, 25]. There is also evidences indicating that this species was mainly piscivorous [6]. Despite these adaptations for active pursuing, *Spinosaurus* probably was a shoreline generalist [26]. This is in line with some observations that spinosaurids were not strictly piscivorous [20, 25]. There is, for instance, evidence of hunting or scavenging of pterosaurs [27]. Probably for these specializations to catch fish and also having a broad diet, *Spinosaurus* was highly abundant in comparison to other top predators in some sites [28].

#### 2.1.3 Other Theropods

In addition to *Spinosaurus*, the Kem Kem beds have at least two other large predators, *Carcharodontosaurus saharicus* and *Deltadromeus agilis* [1]. These large-bodied predators were probably opportunistic scavengers, as evidenced for large Theropods in general [16, 29, 30]. In the present work, I consider that *Spinosaurus* can interact with *Carcharodontosaurus* and/or *Deltadromeus*. For the sake of simplicity, I consider these latter two to belong to the functional group of large land predators, and in the context of this work I call this group “non-*Spinosaurus* Theropods” (NST, hereinafter).

### 2.2 Models

Krivan and Schmitz [14] proposed a model with DMI in which the density of one species exerts a reduction in the foraging activity of the other species. Furthermore, O’Bryan et al. [31] proposed a model for carrion-scavenger dynamics. These two models are general, meaning that they can be applied to a vast class of biological systems that share the assumptions of each model. Taking this into consideration, based on the building blocks from [14] and [31], I propose the following model:

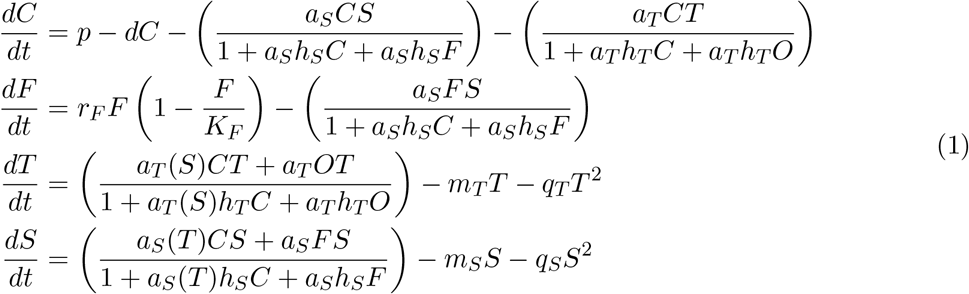

in which the four compartments are Carrion *C*, Fishes *F*, NST *T*, and *Spinosaurus S*. The generation of carrion biomass *p* is due to terrestrial animal death, and *d* is the rate of decomposition. I consider the following model components: a Holling type-II multispecies functional response [32, 33] for both top predators, in which *a*_*S*_ and *a*_*T*_ are the attacking rates of *Spinosaurus* and the NST, respectively, with *h*_*S*_ and *h*_*T*_ their manipulating time; a logistic growth for the fish community with *r*_*F*_ the intrinsic growth rate and *K*_*F*_ is the carrying capacity of the fish community; a linear density-independent *m*_*S*_ and *m*_*T*_ for *Spinosaurus* and the NST, respectively; and also a quadratic density-dependent mortality *q*_*S*_ and *q*_*T*_ for each top predator. This density-dependent mortality can mean, for example, effects of intraspecific competition or cannibalism [34]. The functions *a*_*T*_ (*S*) and *a*_*S*_(*T*) can assume two forms, representing two distinct scenarios: resource competition with DMIs, and purely resource competition. They are displayed in Table 1. The parameters *λ*_*S*_ and *λ*_*T*_ are the attacking coefficients of *Spinosaurus* and NST in the absence of its competitor, respectively; and *α* is the intensity of the DMI that *Spinosaurus* exerts on NST, and *β* is the NST intensity over *Spinosaurus*. The results concerning the scenario of no DMI are presented in Supplementary Material A.

**Table 1:** Considered mathematical functions for *a*_*T*_ (*S*) and *a*_*S*_(*T*) in the present work. Two scenarios are evaluated: the occurrence of DMI and the lack of DMI.

| Type | Equations |
| --- | --- |
| DMI | $a_T(S) = \lambda_T \exp(-\alpha S)$<br>$a_S(T) = \lambda_S \exp(-\beta T)$ |
| No DMI | $a_T(S) = a_T$<br>$a_S(T) = a_S$ |

### 2.3 Model settings

Regarding the parameter values, they were chosen to yield coexistence in the two studied scenarios. To understand the effects of the parameters, I employed numerical continuation of parameter [35, 36] to understand the influence of two parameters, *r*_*F*_ and *O*, on the system. The increase of *r*_*F*_ means enrichment of resources in the aquatic environment, and the increase of *O* means an increase in the availability of terrestrial food sources. In addition, the numerical continuation of *p* is presented in Supplementary Material B.

I also analysed the extreme case in which one or more food sources are not available for the top predators. With this purpose, I simulated four cases: (i) the terrestrial herbivores are absent; (ii) the fish community is absent; (iii) both terrestrial and aquatic food sources are absent; (iv) variation of the carrion availability, i.e., the shared resource. These results are presented in Supplementary Material B. In order to complement the study, I also employed a Sensitivity Analysis of parameter [37, 38], which is presented in the Supplementary Material C.

The ordinary differential equations are solved using the LSODA method [39] from SciPy [40]. The Jupyter notebook code to solve the models is available at https://github.com/Tungdil01/palaeoEcologyKemKem.

## 3 Results

A simulated time series of model (1) with top predator’s DMI is presented in Figure 2. In this hypothetical scenario, all species / functional groups coexist in a stable equilibrium dynamics. A similar result is observed for the no DMI scenario, shown in Supplementary Material A. However, the two competitors equilibrium densities are higher with no DMI.

**Figure 2:**
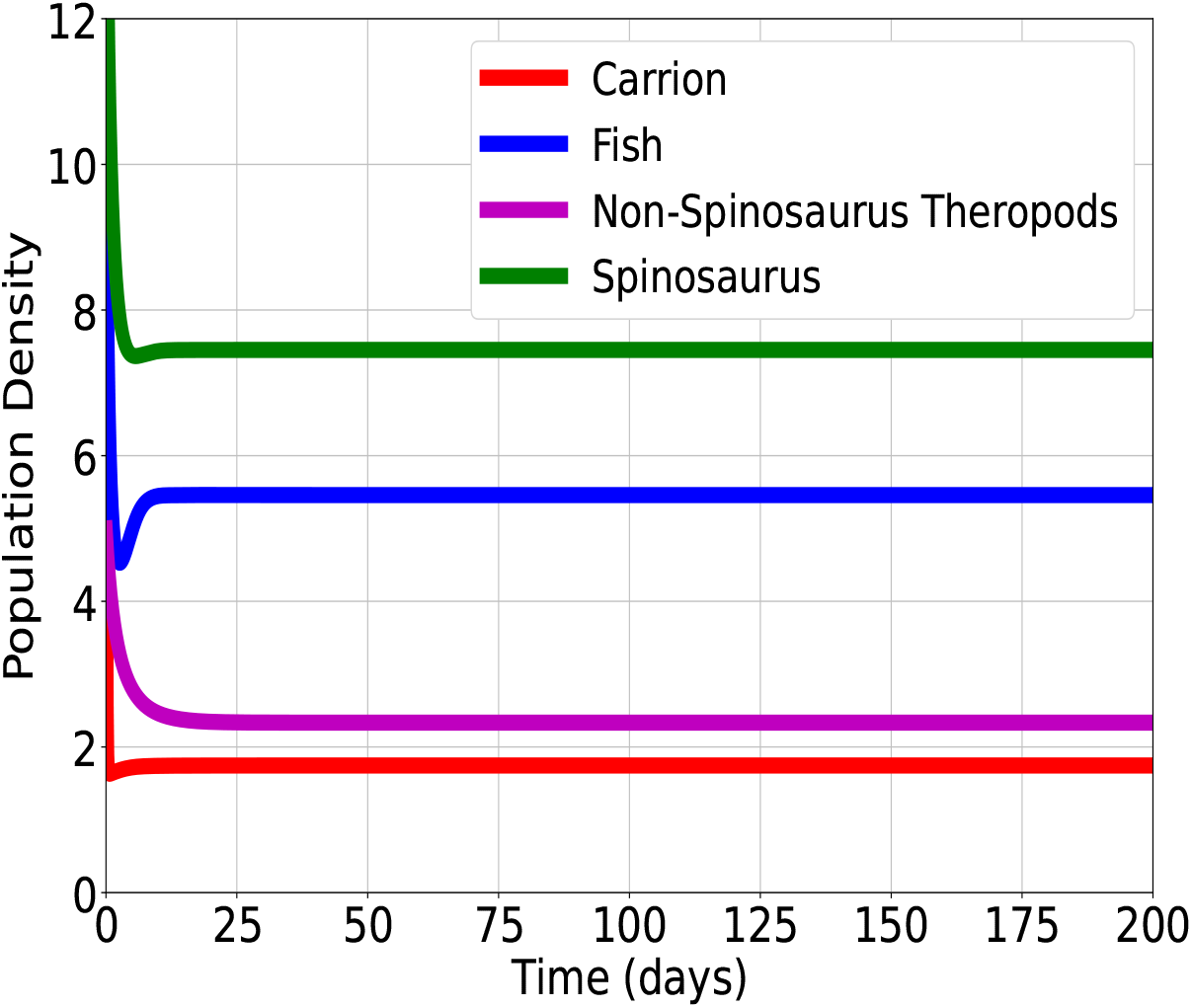
Time series of model (1) including the DMI. The initial conditions of the four compartments are: Carrion *C*(0) = 10, Fishes *F* (0) = 10, NST *T* (0) = 5, *Spinosaurus S*(0) = 15. Parameter values: *λ*_*T*_ = 0.1, *λ*_*S*_ = 0.01, *α* = 1, *β* = 100, *p* = 20, *d* = 10, *a*_*S*_ = 1, *h*_*S*_ = 1, *a*_*T*_ = 1, *h*_*T*_ = 1, *O* = 0.5, *r*_*F*_ = 2, *K*_*F*_ = 10, *m*_*T*_ = 0.1, *q*_*T*_ = 0.1, *m*_*S*_ = 0.1, *q*_*S*_ = 0.1.

Increasing the parameters related to the prey items of each competitor yield an increase in their corresponding densities, which is displayed in Figure 3. Figure 3(a) shows that the increase in *r*_*F*_ increases the equilibrium density of *Spinosaurus* and decreases density of the NST. This decrease in the NST equilibrium density might be a consequence of the DMI, since *β* is much higher than *α*. On the other hand, Figure 3(b) shows an increase in the equilibrium density of the NST with the increase in its food sources availability. The equilibrium density of *Spinosaurus* first slightly decreases (in the order of 1*e* − 02) for small *O*, but then slightly increases for approximately *O* ≥ 2.

**Figure 3:**
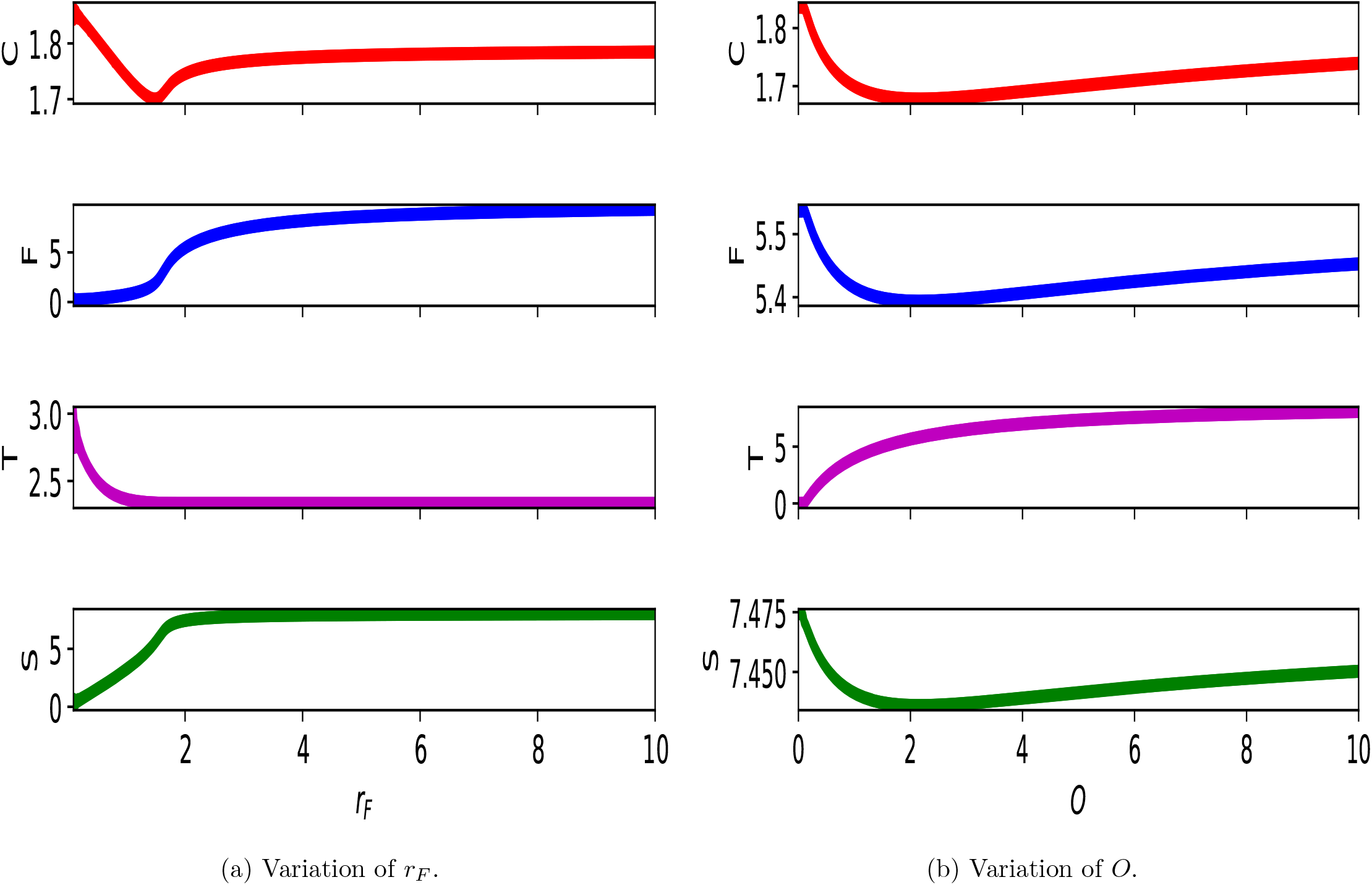
Numerical continuation for model (1) including the DMI. The initial conditions and parameter values are the same as in Figure 2, but in (a) *r*_*F*_ is varied and in (b) *O* is varied. Initial conditions: *C*(0) = 10, *F* (0) = 10, *T* (0) = 5, *S*(0) = 15. Parameter values: *λ*_*T*_ = 0.1, *λ*_*S*_ = 0.01, *α* = 1, *β* = 100, *p* = 20, *d* = 10, *a*_*S*_ = 1, *h*_*S*_ = 1, *a*_*T*_ = 1, *h*_*T*_ = 1, *O* = 0.5, *r*_*F*_ = 2, *K*_*F*_ = 10, *m*_*T*_ = 0.1, *q*_*T*_ = 0.1, *m*_*S*_ = 0.1, *q*_*S*_ = 0.1.

## 4 Discussion

The present paper intended to study a simple ecological model, representing a piece of the Kem Kem Group, and to construct a numerical framework to analyse this system. The results presented in this work suggest that the answer to the Stomer’s riddle is niche partitioning. This could have allowed then the Kem Kem palaeoenvironment to sustain a relatively high number of large predators in comparison to large herbivores, provided there are sufficient food sources. As displayed in Figure 3, the densities of the two top predators are highly dependent on the availability of exclusive food sources, i.e., fishes for *Spinosaurus* and sauropods and others for the NST (see also Supplementary Material B). As far as carrion is concerned, the simulations suggest that it alone may not be able to sustain the large predators guild for the scenario of DMI (Supplementary Material B). Moreover, as carrion is increased, the fish community also increases (Supplementary Material B). This result may be due to the phenomenon of predator satiation [41], which in the case of predation by *Spinosaurus* releases the fish community.

Beevor et al. [28] showed evidence supporting a high abundance of *Spinosaurus* teeth in comparison with other Theropods in some sites in Morocco. We could qualitatively relate this information with the results of Figure 3, for the reason that a high aquatic enrichment yields a high equilibrium density of *Spinosaurus*. On the other hand, if the terrestrial herbivore sources are kept constant, the NST equilibrium density is also kept at a constant value.

Ecological models such as those developed in this paper often describe a range of extant biological systems, like terrestrial mammals [42], small invertebrates [43], and marine communities [44], just to cite a few examples. Also concerning ecological models, Pahl and Ruedas [16] employed an agent-based technique to study a system in which carnosaurs are scavengers. They showed that sauropod carrion could sustain several individuals and scavenging could be quite common in large Theropods, which could explain few predatory specializations in carnosaurs. This could justify the fact that the NST survived even in the extreme case of the absence of other food sources (Supplementary Material B).

An interchange between palaeoecology and ecological modelling can be further explored and deepened for other palaeoenvironments, with the possibility to provide palaeontologists potential ecological mechanisms in the world today to explain some patterns that probably also occurred in the past. Concerning the limitations of the present analysis, it is important to note that a simplified biological setup was employed. An actual ecological network is composed of many elements, some of which were neglected in the modelling process. Future analyses can detail the ecological network by including more species in the food web. Some potential extensions are: (i) the inclusion of more than two competing large predators; (ii) the decoupling of the “fishes” compartment to spotlight individual species; and (iii) the examination of terrestrial herbivore dynamics.

## 5 Conclusions

The density-mediated interaction (DMI) was evaluated by the modelling framework developed in this work. This type of interaction reduced the equilibrium densities of the two competitors in comparison with the no DMI scenario. One large predator might even go extinct in extreme cases in the absence of a food source. Another aspect regarding the variation in the food sources is that in the scenario of DMI, increasing the food source of a predator causes a decrease in the density of its competitor. As a consequence of further increasing the food source, the other predator stays with a fixed density or has its density slightly increased. Finally, increasing the carrion availability, *Spinosaurus* and fish densities increased, but the non-*Spinosaurus* Theropods (NST) had a constant density in the DMI scenario, indicating that the intensity of DMI can have a major influence on the dynamical outcomes.

## Acknowledgements

I would like to thank Diego T. Volpatto and Felipe F. Barbosa for their kind and supportive help with the code and the scientific nomenclature, respectively. This work was supported by a fellowship from the Institutional Training Program (PCI) from the Ministry of Science, Technology, Innovation and Communication (MCTIC) (Grant Number: 301327/2020-3).

