## Supplementary Material for "Population modelling insights of extinct environments: the case of the Kem Kem palaeocommunity"

### A Results for the scenario of no density-mediated interaction

The model studied in the main text includes a density-mediated interaction (DMI), as shown below:

$$\begin{aligned}
\frac{dC}{dt} &= p - dC - \left(\frac{a_S CS}{1 + a_S h_S C + a_S h_S F}\right) - \left(\frac{a_T CT}{1 + a_T h_T C + a_T h_T O}\right) \\
\frac{dF}{dt} &= r_F F \left(1 - \frac{F}{K_F}\right) - \left(\frac{a_S FS}{1 + a_S h_S C + a_S h_S F}\right) \\
\frac{dT}{dt} &= \left(\frac{a_T(S)CT + a_T OT}{1 + a_T(S) h_T C + a_T h_T O}\right) - m_T T - q_T T^2 \\
\frac{dS}{dt} &= \left(\frac{a_S(T)CS + a_S FS}{1 + a_S(T) h_S C + a_S h_S F}\right) - m_S S - q_S S^2
\end{aligned} \tag{1}$$

with $a_T(S) = \lambda_T \exp -\alpha S$ and $a_S(T) = \lambda_S \exp -\beta T$.

However, in this section I consider the scenario with no DMI, i.e., $a_T(S) = a_T$ and $a_S(T) = a_S$. Therefore, the model reads:

$$
\begin{aligned}
\frac{dC}{dt} &= p - dC - \left(\frac{a_S CS}{1 + a_S h_S C + a_S h_S F}\right) - \left(\frac{a_T CT}{1 + a_T h_T C + a_T h_T O}\right) \\
\frac{dF}{dt} &= r_F F \left(1 - \frac{F}{K_F}\right) - \left(\frac{a_S FS}{1 + a_S h_S C + a_S h_S F}\right) \\
\frac{dT}{dt} &= \left(\frac{a_T CT + a_T OT}{1 + a_T h_T C + a_T h_T O}\right) - m_T T - q_T T^2 \\
\frac{dS}{dt} &= \left(\frac{a_S CS + a_S FS}{1 + a_S h_S C + a_S h_S F}\right) - m_S S - q_S S^2
\end{aligned}
\tag{A.1}
$$

The time series and the numerical continuation are presented in Figures A.1 and A.2, respectively. The results are similar to those presented in the main text, with the difference that when there is no DMI, the top predators have higher equilibrium densities compared with the scenario in which the DMI occurs.

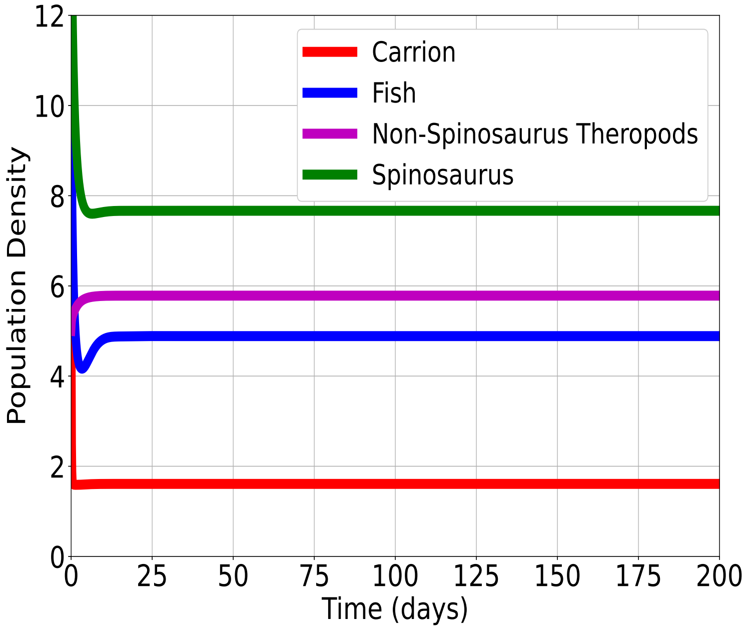


Figure A.1: Time series of model (A.1). The initial conditions and parameter values are the same as in the main text. The initial conditions of the four compartments are: Carrion $C(0) = 10$, Fishes $F(0) = 10$, non-*Spinosaurus* Theropods (NST) $T(0) = 5$, *Spinosaurus* $S(0) = 15$. Parameter values: $p = 20, d = 10, a_S = 1, h_S = 1, a_T = 1, h_T = 1, O = 0.5, r_F = 2, K_F = 10, m_T = 0.1, q_T = 0.1, m_S = 0.1, q_S = 0.1$.

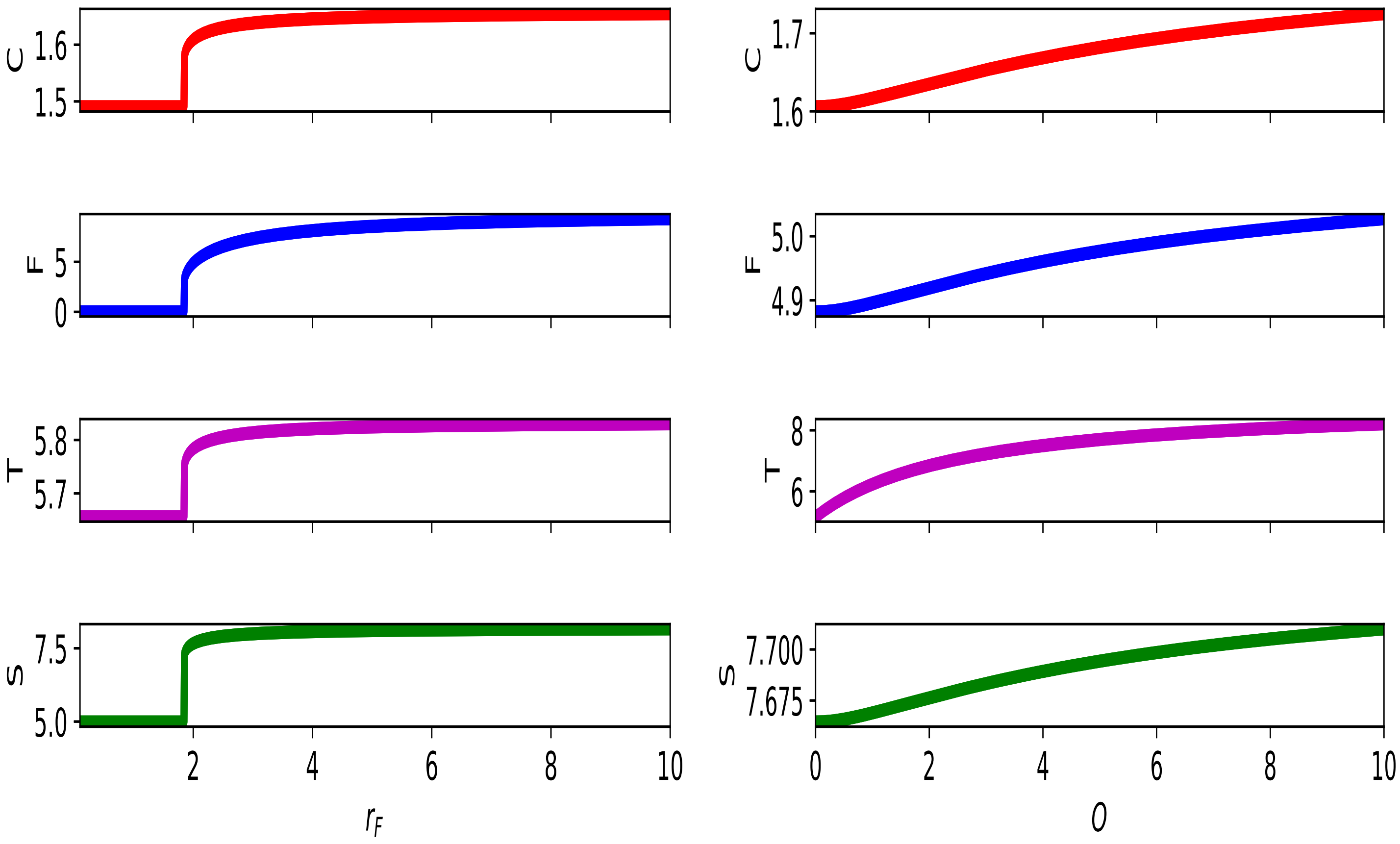


(a) Variation of $r_F$.

(b) Variation of $O$.

Figure A.2: Numerical continuation for model (A.1). The initial conditions and parameter values are the same as in Figure A.1, but in (a) $r_F$ is varied and in (b) $O$ is varied. Initial conditions: $C(0) = 10$, $F(0) = 10$, $T(0) = 5$, $S(0) = 15$. Parameter values: $p = 20, d = 10, a_S = 1, h_S = 1, a_T = 1, h_T = 1, O = 0.5, r_F = 2, K_F = 10, m_T = 0.1, q_T = 0.1, m_S = 0.1, q_S = 0.1$.

### B Study of food restriction

In this section, I study the consequences of restricting one or both food sources for the two top predators, considering the model with DMI (1) and with no DMI (A.1).

#### B.1 Absence of exclusive food sources

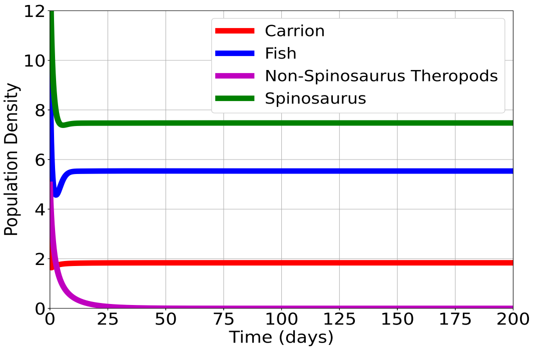


(a) DMI with absence of $O$.

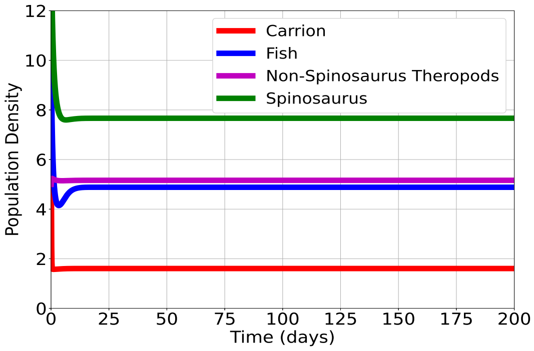


(b) No DMI with absence of $O$.

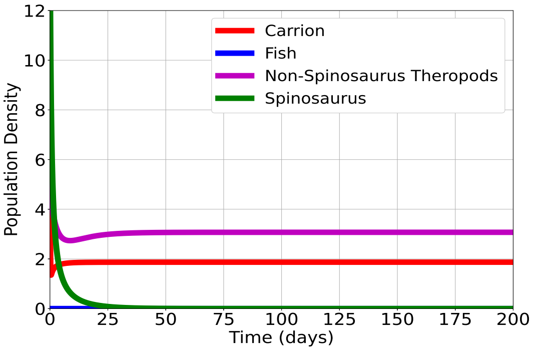


(c) DMI with absence of Fishes $F$.

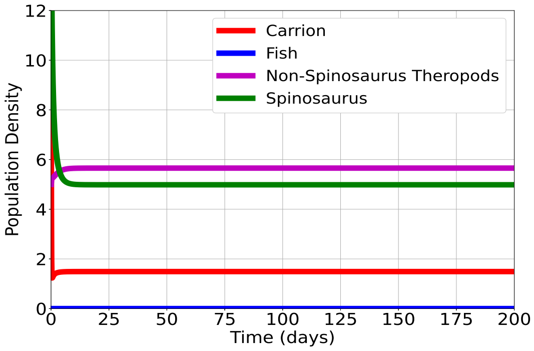


(d) No DMI with absence of Fishes $F$.

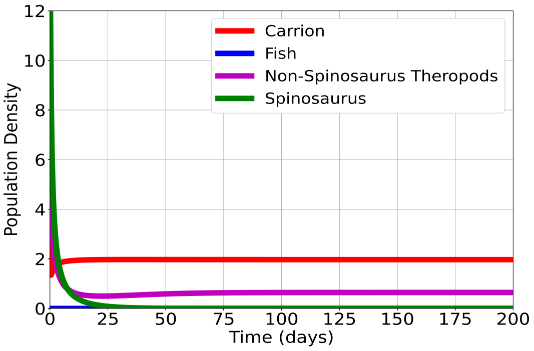


(e) DMI with absence of $F$ and $O$.

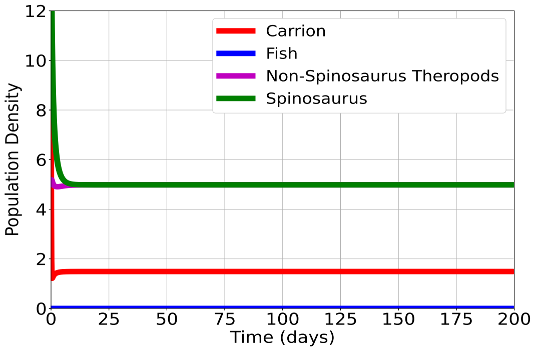


(f) No DMI with absence of $F$ and $O$.

Figure B.1: Scenarios with the absence of a given food source.

The initial conditions and parameter values of Figure B.1 are the same as in Figure A.1, with the addition of $\lambda_T = 0.1, \lambda_S = 0.01, \alpha = 1, \beta = 100$ for the scenario with DMI given by model (1). Figure B.1(a,c,e) shows the outcomes for the absence of a food source for the scenario in which occurs the DMI. In the absence of $O$, the NST is locally extinct in the environment. In the absence of Fishes $F$, *Spinosaurus* who becomes locally extinct. For the employed parameter values, when $O$ and $F$ are absent, the *Spinosaurus* is locally extinct.

The results when there is no DMI are displayed in Figure B.1(b,d,f). Contrarily to the DMI scenario, the no DMI scenario does not show local extinction for the three combinations of food source absence. This means that the Carrion can sustain the two top predator populations even in the absence of exclusive food sources.

#### B.2 Variation of Carrion (shared resource)

Figure B.2 shows the different outcomes of increasing the parameter $p$ for model (1) and (A.1). Figure B.2(a) shows that the Carrion and Fishes increase, while the predators remain with constant densities.

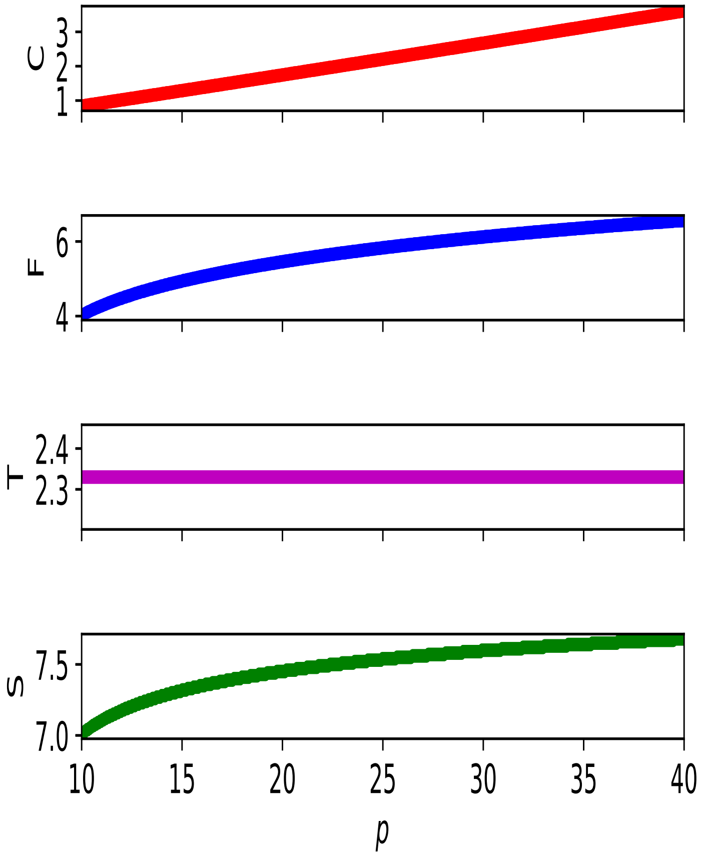


(a) Variation of $p$ for model (1).

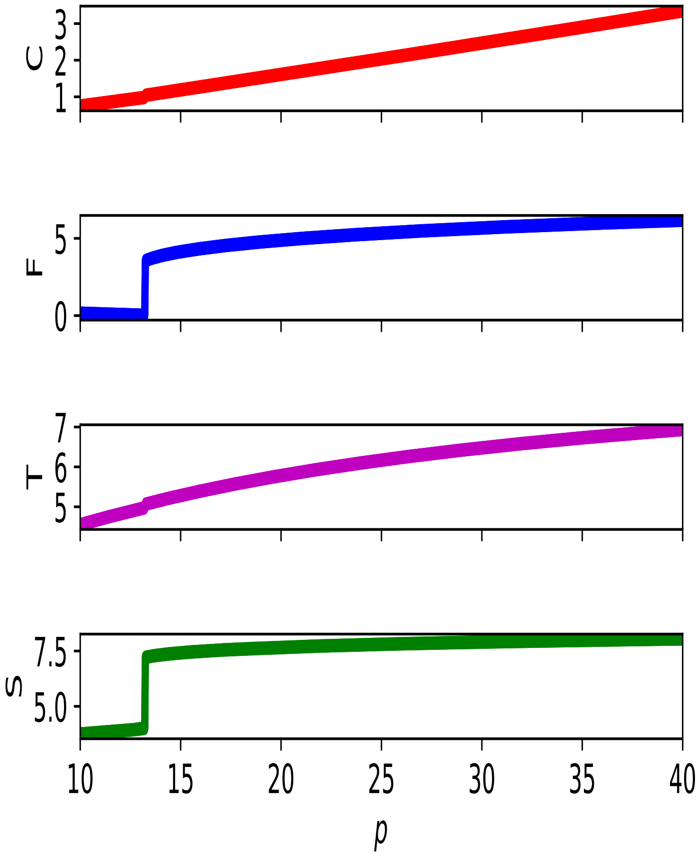


(b) Variation of $p$ for model (A.1).

Figure B.2: Numerical continuation for (a) model (1) including the DMI, and (b) model (A.1) with no DMI. Initial conditions: $C(0) = 10$, $F(0) = 10$, $T(0) = 5$, $S(0) = 15$. Parameter values: $\beta = 100, p = 20, d = 10, a_S = 1, h_S = 1, a_T = 1, h_T = 1, O = 0.5, r_F = 2, K_F = 10, m_T = 0.1, q_T = 0.1, m_S = 0.1, q_S = 0.1$ for both scenarios, and $\lambda_T = 0.1, \lambda_S = 0.01, \alpha = 1$ for the scenario with the DMI.

### C Sensitivity Analyses

I employ the Morris Elementary Effects Method [1, 2] in the present paper. This is a global sensitivity screening method that provides two sensitivity indexes, $\mu_i^*$ and $\sigma_i, i = 1, \ldots, d$, regarding a quantity of interest (QoI), here assumed to be the density of each top predator. In the latter expression, $d$ is the total number of uncertain parameters. The average $\mu^*$ gives the relative importance of each parameter concerning the QoI and the standard deviation $\sigma$ describes their non-linear effects. For more details on the method, see [2, 3]. The method was implemented using the SALib package [4].

In the rest of this section, I show the results of the Sensitivity Analysis of the three studied models. We can observe the evolution of the importance of each parameter according to the two indexes, $\mu^*$ and $\sigma$. These indexes indicate the most important parameters for the given QoI, and doing so, can reveal which terms of the models are dominating. The initial conditions and parameter values are the same as in Figure A.1, with the addition of $\lambda_T = 0.1, \lambda_S = 0.01, \alpha = 1, \beta = 100$ for the scenario with DMI given by model (1).

#### C.1 DMI

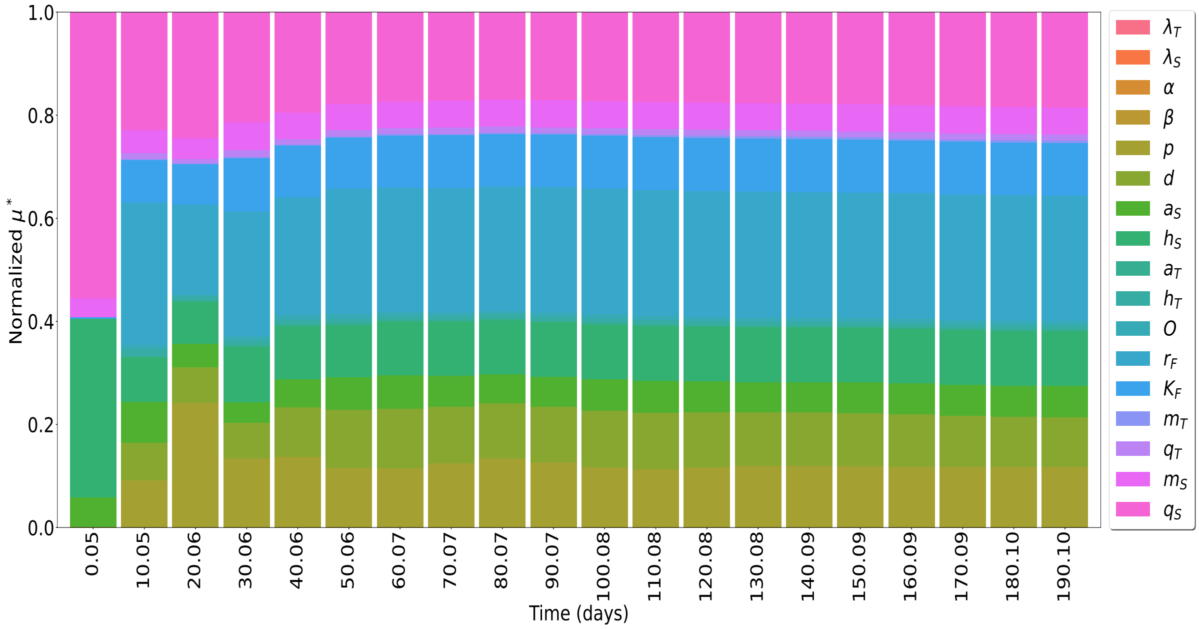


(a) $\mu^*$.

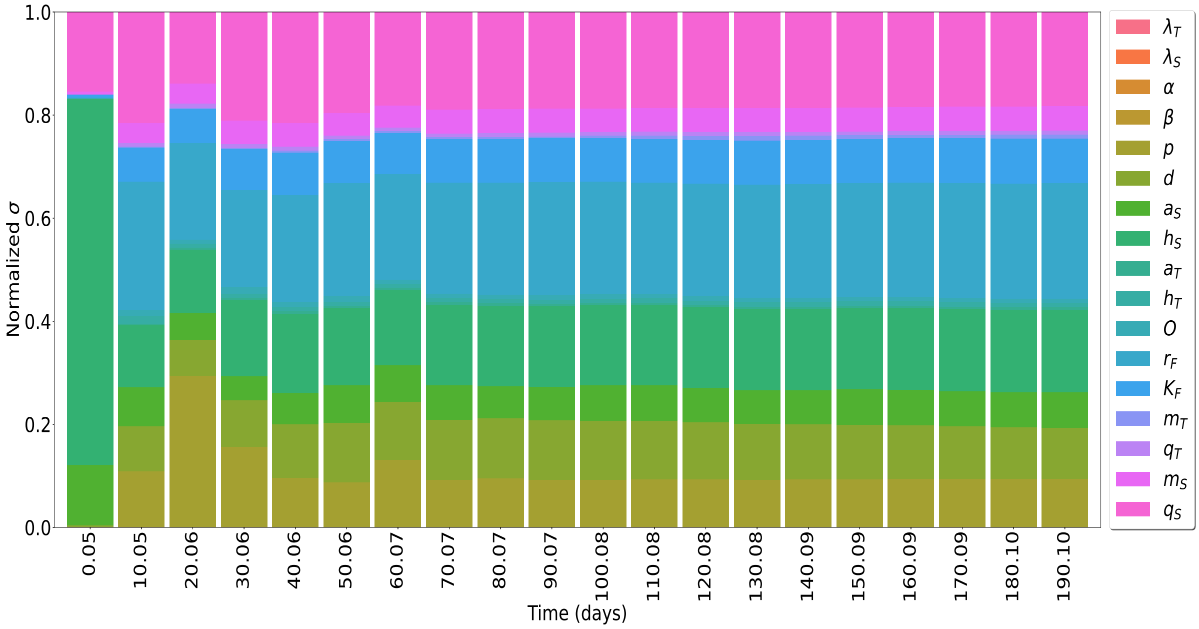


(b) $\sigma$.

Figure C.1: Sensitivity analysis using the density of *Spinosaurus* as the QoI. The five most important parameters are (a) $\mu^*$: $r_F, q_S, p, h_S, K_F$; and (b) $\sigma$: $r_F, q_S, h_S, d, p$.

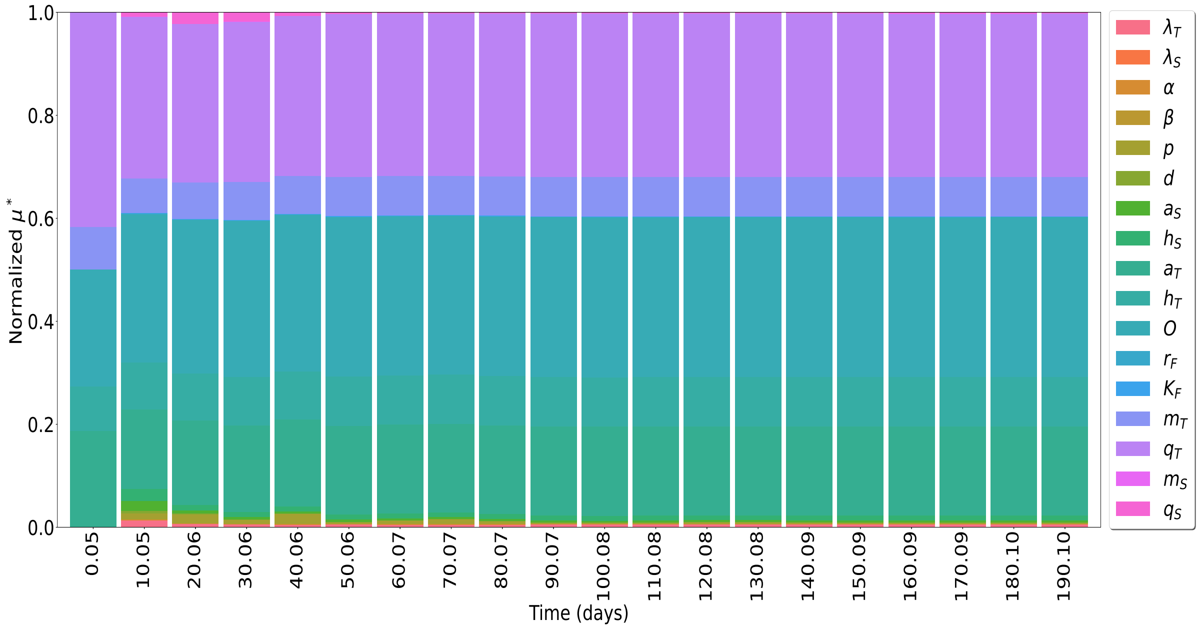


(a) $\mu^*$.

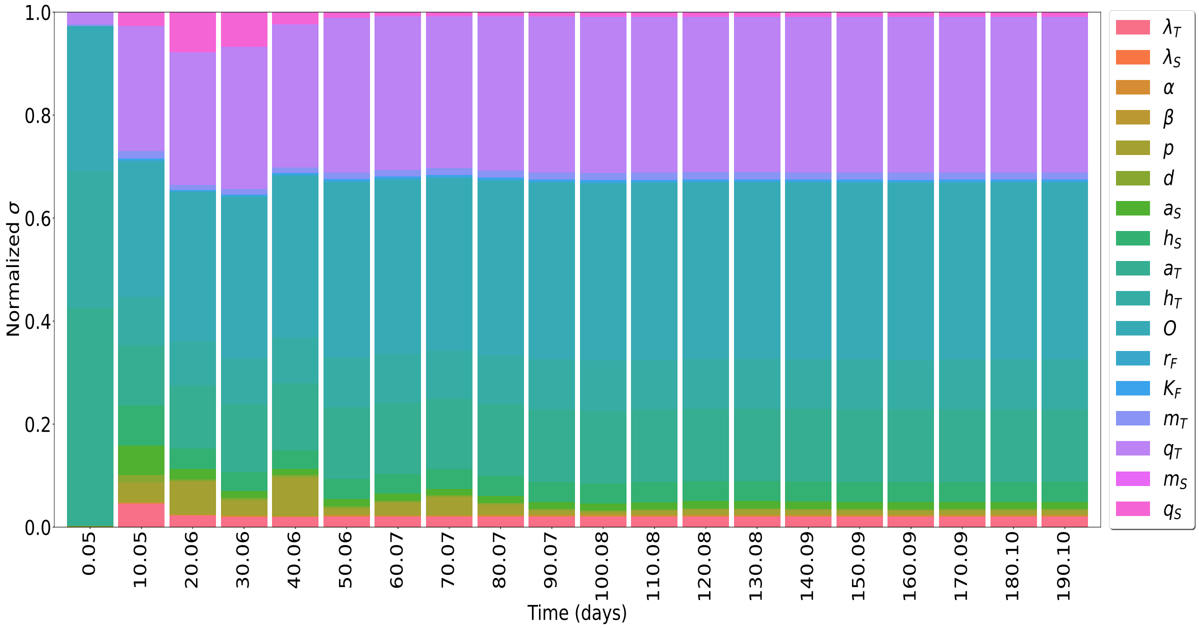


(b) $\sigma$.

Figure C.2: Sensitivity analysis using the density of the NST as the QoI. The five most important parameters are (a) $\mu^*$: $q_T, O, a_T, h_T, m_T$; and (b) $\sigma$: $O, q_T, a_T, h_T, h_S$.

#### C.2 No DMI

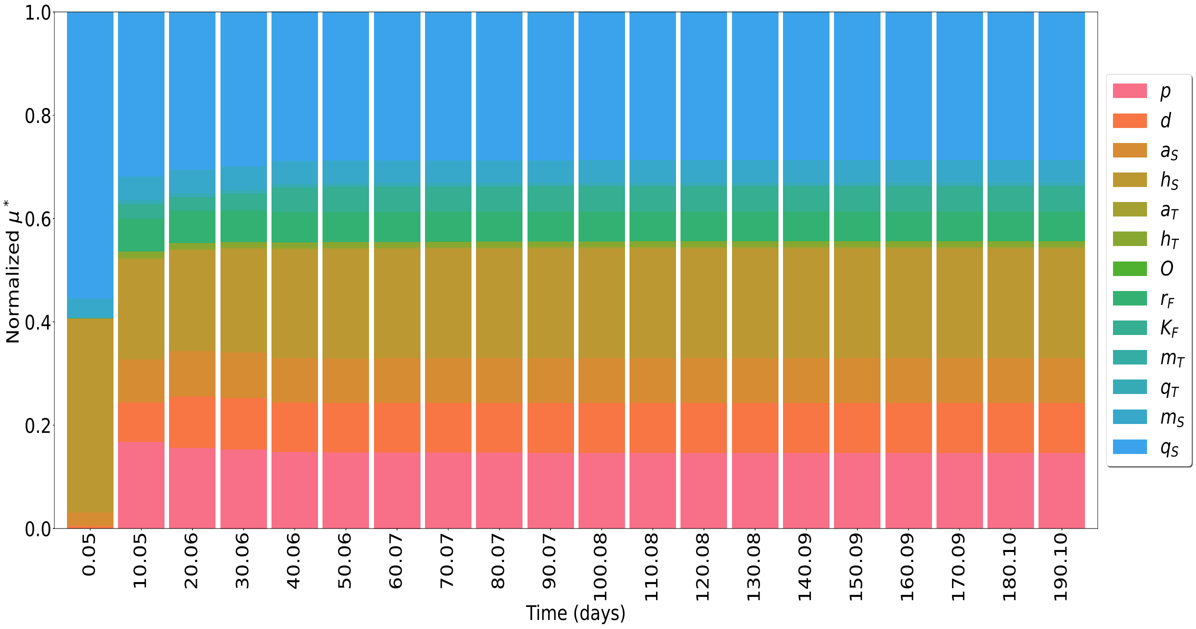


(a) $\mu^*$.

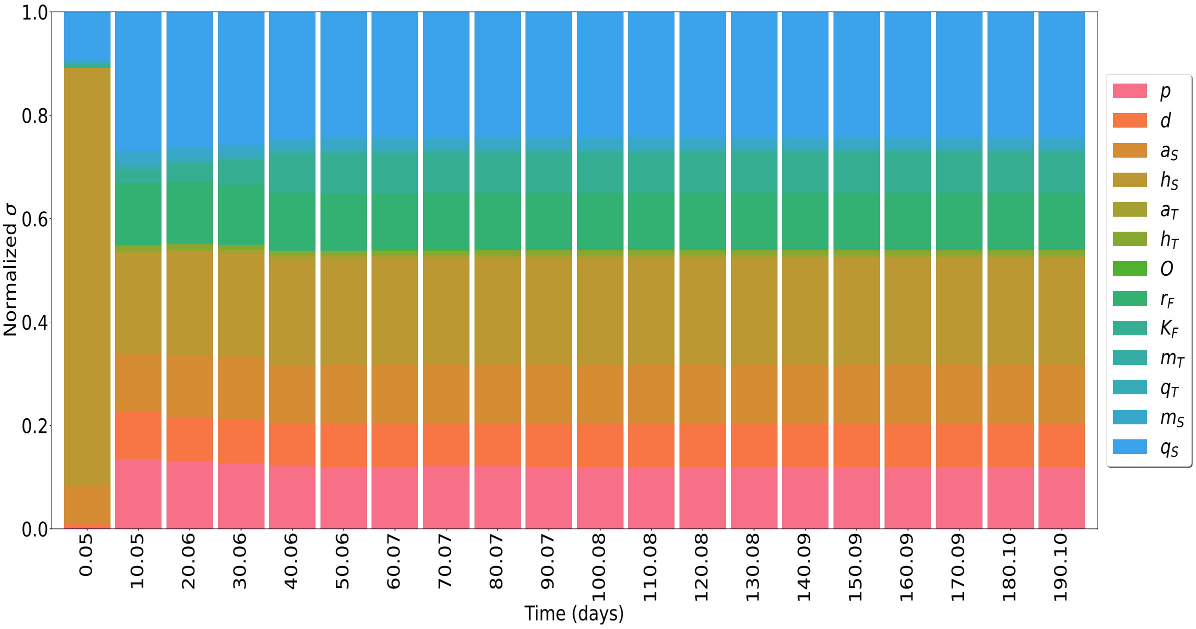


(b) $\sigma$.

Figure C.3: Sensitivity analysis using the density of the NST as the QoI. The five most important parameters are (a) $\mu^*$: $q_S, h_S, p, a_S, d$; and (b) $\sigma$: $h_S, q_S, p, d, a_S$.

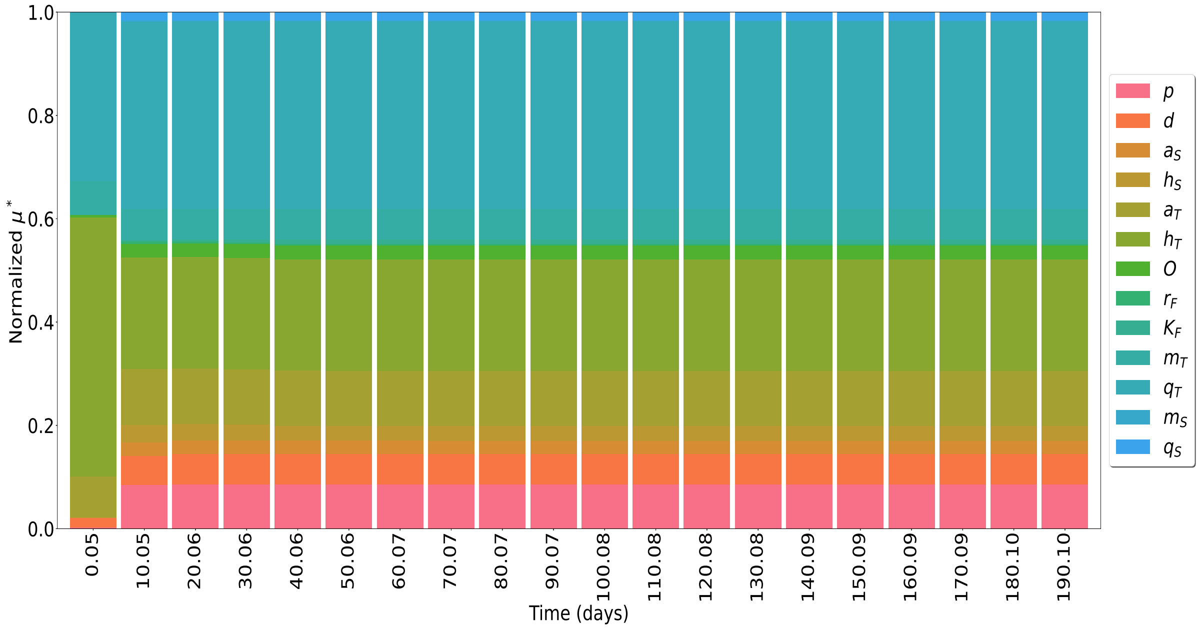


(a) $\mu^*$.

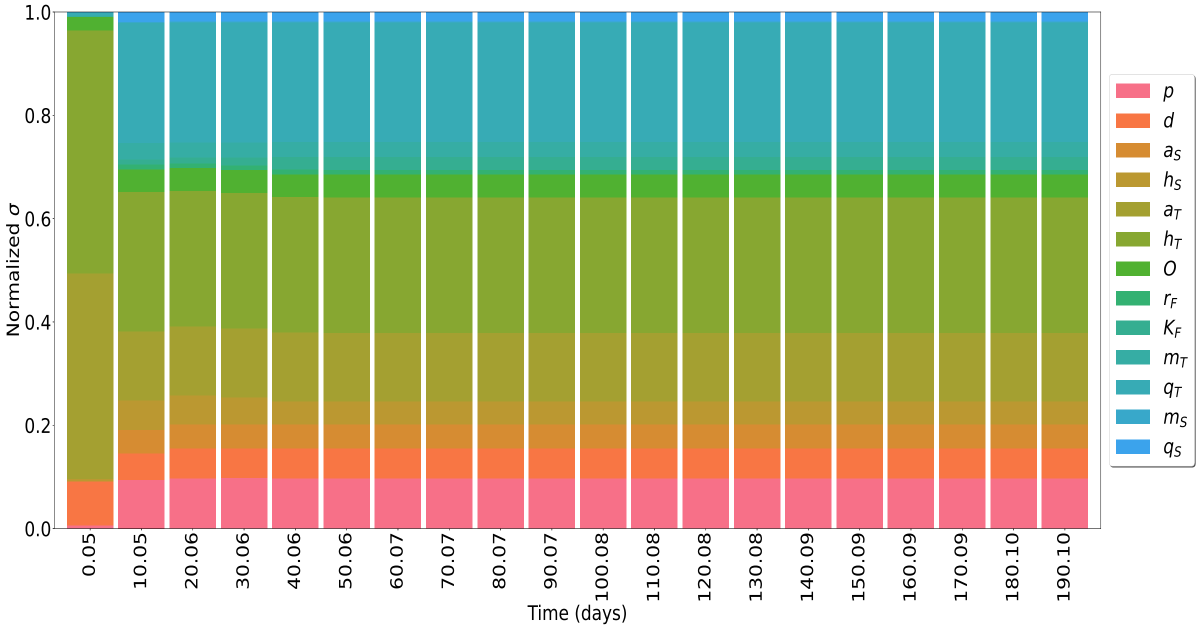


(b) $\sigma$.

Figure C.4: Sensitivity analysis using the density of the NST as the QoI. The five most important parameters are (a) $\mu^*$: $q_T, h_T, a_T, p, m_T$; and (b) $\sigma$: $h_T, q_T, a_T, p, a_S$.
